# Spatiospectral signatures of stomach-brain synchrony

**DOI:** 10.64898/2026.03.13.711584

**Authors:** Teresa Berther, Martina Saltafossi, Jana Fehring, Ignacio Rebollo, Daniel S. Kluger

## Abstract

While cognitive neuroscience has increasingly recognized the brain-body connection, the temporal dynamics of how the stomach’s intrinsic rhythm (∼0.05 Hz) modulates human neural activity remain poorly understood. Leveraging high-resolution magneto-encephalography (MEG) and novel high-density electrogastrography (EGG), we provide the first comprehensive spatiotemporal mapping of human gastric-brain coupling at rest. Our circular-linear correlation approach revealed widespread, broadband phase-amplitude coupling between the gastric rhythm and spontaneous cortical oscillations across delta, theta, alpha, and beta bands. Using nonnegative matrix factorization, we identified distinct spatiospectral fingerprints that significantly overlap with the known fMRI-based gastric resting-state network. Functional connectivity analyses based on multivariate Granger causality further showed that these modulation patterns extend to a gastric influence on frequency-specific, directed communication between network components. Crucially, we discovered a consistent preferred gastric phase for these modulations, i.e. during the transition between stomach waves, that was stable across brain regions, frequencies, connections, and time. These findings suggest that the gastric rhythm – potentially in unison with other physiological signals – provides a stable, global scaffold for large-scale oscillatory brain organization. By unravelling these spectral signatures, our results establish the stomach as a critical driver of the temporal coordination of human brain dynamics.

## Introduction

In recent years, cognitive neuroscience has shifted away from its perception of the brain as an isolated, closed-system: Instead, it has increasingly acknowledged that the brain constantly receives input from visceral organs. Vice versa, these visceral inputs are now being recognized as important drivers of brain activity^1^. Variations in resting state neural activity may reflect the sensing and integration of visceral stimuli – a process commonly described as interoception^2^. Besides the bi-directional interplay between the brain and interoceptive signals regulating bodily needs, some visceral organs like the heart, lungs, and stomach also create slow frequency rhythms which directly contribute to the temporal organisation of brain dynamics^1,3^. While the influence of cardiorespiratory rhythms has been widely discussed^4–6^, literature on visceral oscillations within the gastrointestinal system, particularly stomach-related modulation of neural activity, is scarce.

This oscillatory activity in the stomach is intrinsically generated by the specialised interstitial cells of Cajal at the interface between the enteric nervous system and the gastric smooth muscles in the form of electrical slow waves at a frequency of ∼ 0.05 Hz (corresponding to a cycle length of approx. 20s). The gastric rhythm sets the pace of muscle contractions during digestion, but is generated at all times^7^, even in the absence of contractions or if the stomach is decoupled from the brain^8^.

The stomach poses an interesting focus as an external oscillator possibly influencing large-scale brain organisation for several reasons. First, visceral signals from the stomach are projected to subcortical and cortical targets including amygdala, thalamus, primary and secondary somatosensory cortices, the insula, the ventromedial prefrontal cortex (vmPFC), and cingulate motor regions, engaging all major neurotransmitter systems^9,10^. Second, Rebollo *et al.*^11^ identified a delayed connectivity resting-state network (RSN) of brain regions whose spontaneous fluctuations in BOLD-activity were modulated by the phase of the gastric rhythm. This network cuts across classical RSNs and partially overlaps with autonomic regulation areas. A follow-up characterization revealed that the gastric network is underrepresented in transmodal regions but overrepresented in unimodal sensory-motor regions^12^. Not only areas tasked with the processing of interoceptive information like the insula, but all sensory and motor cortices were shown to be coupled to the phase of the gastric rhythm, including areas responding to touch, vision, audition, and olfaction, pointing toward a domain-general influence of the gastric rhythm on global neural dynamics.

Lastly, the temporal structure of the amplitude of spontaneous brain oscillations at rest is modulated by gastric phase^13^. Phase-amplitude coupling (PAC) has been widely investigated as an important principle of temporal organisation in large-scale brain activity in humans^14^. From a holistic brain-body perspective, and based on the rich anatomical connectivity linking gut and brain, this would entail that the phase of gastric cycles may play a role in coordinating oscillatory synchrony across the brain. In line with this conceptualisation, gastric electrical stimulation drives local field potentials in rats, resulting in widespread transient and phase-locked broadband neural responses to gastric signals^15^. In humans, transfer entropy – a measure of directed information flow similar to Granger causality – also provided evidence for an ascending modulatory influence of the stomach^13^. Note that the analyses of both BOLD-activity as well as phase-amplitude coupling applied to quantify gastric-brain coupling are inherently correlational in nature, thus some caution regarding true causality is warranted.

To conclude, current animal and human literature indicates a widespread gastric modulation of brain activity and provides a precise anatomical mapping of these effects. The temporal dynamics of the modulation however are largely unknown, both on the cortical side – i.e., the time-frequency domain – and on the gastric side. Available evidence is predominantly derived from fMRI studies, a methodology ill-suited for temporal characterisation. To date, only a single study has investigated stomach-brain synchrony with more temporally precise measurements of brain activity using magnetencephalography (MEG), but was restricted to the alpha frequency range (8 – 13 Hz)^13^. Here, we provide the first comprehensive spatiotemporal mapping of human gastric-brain coupling at rest. Leveraging oscillatory analyses of high-resolution MEG and a novel application of high-density electrogastrography (EGG), we replicate the anatomical layout and unravel spectral signatures of stomach-brain dynamics across the frequency spectrum. We first quantified gastric modulation of source-level neural oscillations throughout the whole brain, introducing a novel circular-linear correlation approach for PAC across a wide range of frequencies. Dimensionality reduction in the spatial domain revealed a constrained anatomical map of gastric phase-dependent amplitude modulations. We further identified the specific spectral fingerprint of each of these network components, and finally assessed the temporal consistency of gut-brain interactions. Finally, we characterized gastric modulation of frequency-specific functional connectivity between the coupled nodes using a gastric-phase-resolved application of pairwise multivariate Granger causality (mGC).

Our results not only demonstrate that the precise spatial extent of the human gastric network translates across recording domains, but also – for the first time – characterise gut-brain coupling within this network in the spectral domain, revealing broadband modulation and distinct, region-specific fingerprints of gastric brain activity. Analyses of the temporal dynamics of this modulatory effect further revealed a global pattern in stomach-brain synchrony, suggesting a common underlying mechanism and a role of the gastric rhythm in large-scale brain organisation. Consistent with this proposed role, the same shared temporal pattern was reflected in phase-dependent modulation of directed cortico-cortical communication, indicating a gastric influence beyond local oscillatory activity.

## Results

### A gastric clock for cross-spectral cortical dynamics

To evaluate gastric modulation of oscillatory amplitude across the brain, we first extracted the phase time series of the EGG electrode showing maximum power in the normogastric range (0.033 – 0.066 Hz) for further analysis (*M*_EGGfreq_ = 0.047 Hz, *SD* = 0.004 Hz), following established recommendations from Wolpert *et al.*^16^ (cf. Fig. 1a). Source-level oscillatory activity was reconstructed using a beamformer approach and aggregated into parcel-level time series using a recent, multimodal parcellation of the cerebral cortex^17^. We then obtained time-resolved power spectra for each parcel and related them to gastric phase, yielding phase-resolved power spectra in the frequency range between 0.5 and 30 Hz (see Methods for details).

**Fig 1.**
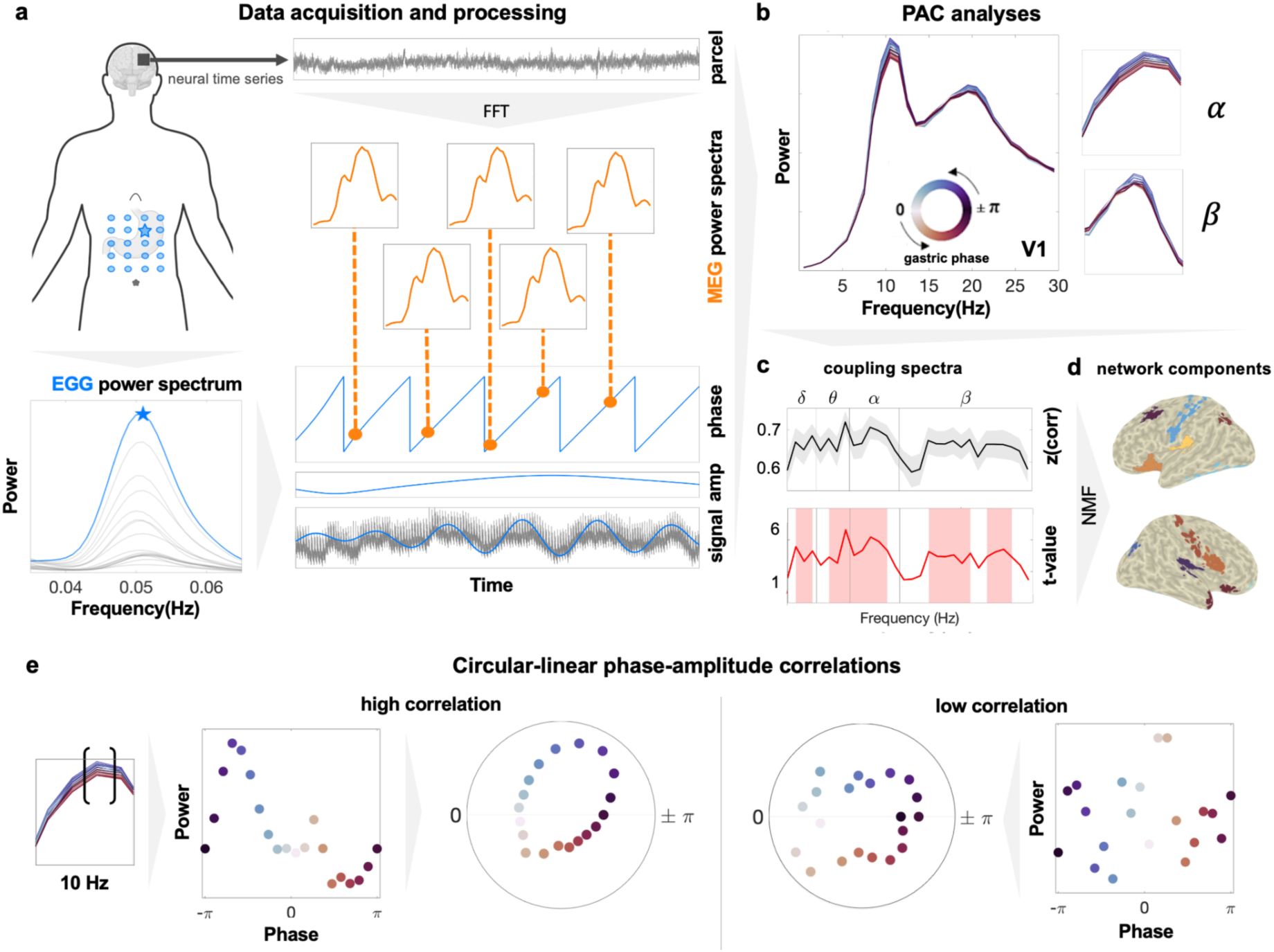
Methods outline. **a**, Data acquisition and processing. EGG was recorded with a high-density electrode layout; bottom spectrogram shows the power spectrum of the electrode with highest power in normogastric range (highlighted). EGG phase and amplitude time series were extracted via Hilbert transform after filtering raw peak electrode signal. Brain activity was recorded with MEG, source reconstructed via beamformer, and aggregated into cortical parcels. Time-resolved power-spectra were computed for each parcel using a moving-window FFT. **b,** Phase-amplitude coupling (PAC) analysis. MEG power spectra were binned into 20 phase bins across the gastric cycle. PAC was computed separately for each frequency and parcel using circular-linear correlation of MEG power and gastric phase. **c,** Exemplary result spectra for parcel V1 in the left hemisphere. Upper panel shows average Fisher-transformed correlation values across the frequency spectrum with SEM shading across participants. Lower panel depicts cluster-permutation t-values with red shading marking frequency ranges showing significant PAC. **d**, Applying nonnegative matrix factorisation to the coupling spectra for dimensionality reduction yielded a spatially constrained map of components with specific spectral profiles of gastric-brain coupling. **e**, Circular-linear correlations. Phase-binned MEG power across gastric phases in linear and circular representation for exemplary frequency of 10 Hz. Left panel shows representation of a high circular-linear correlation between MEG power (amplitude) and gastric phase for the exemplary parcel V1 at 10 Hz, right panel shows low correlation for some simulated exemplary data, respectively.

PAC was determined on the single-subject level as the circular-linear correlation of power and gastric phase for each frequency and parcel separately (see Fig. 2a). Comparison with chance-level coupling correlation values using cluster-based permutation testing revealed widespread gastric modulation of spontaneous brain activity across the whole cortex and frequency spectrum (cf. effect spectra and matrix in Fig. 2a and b).

**Fig 2.**
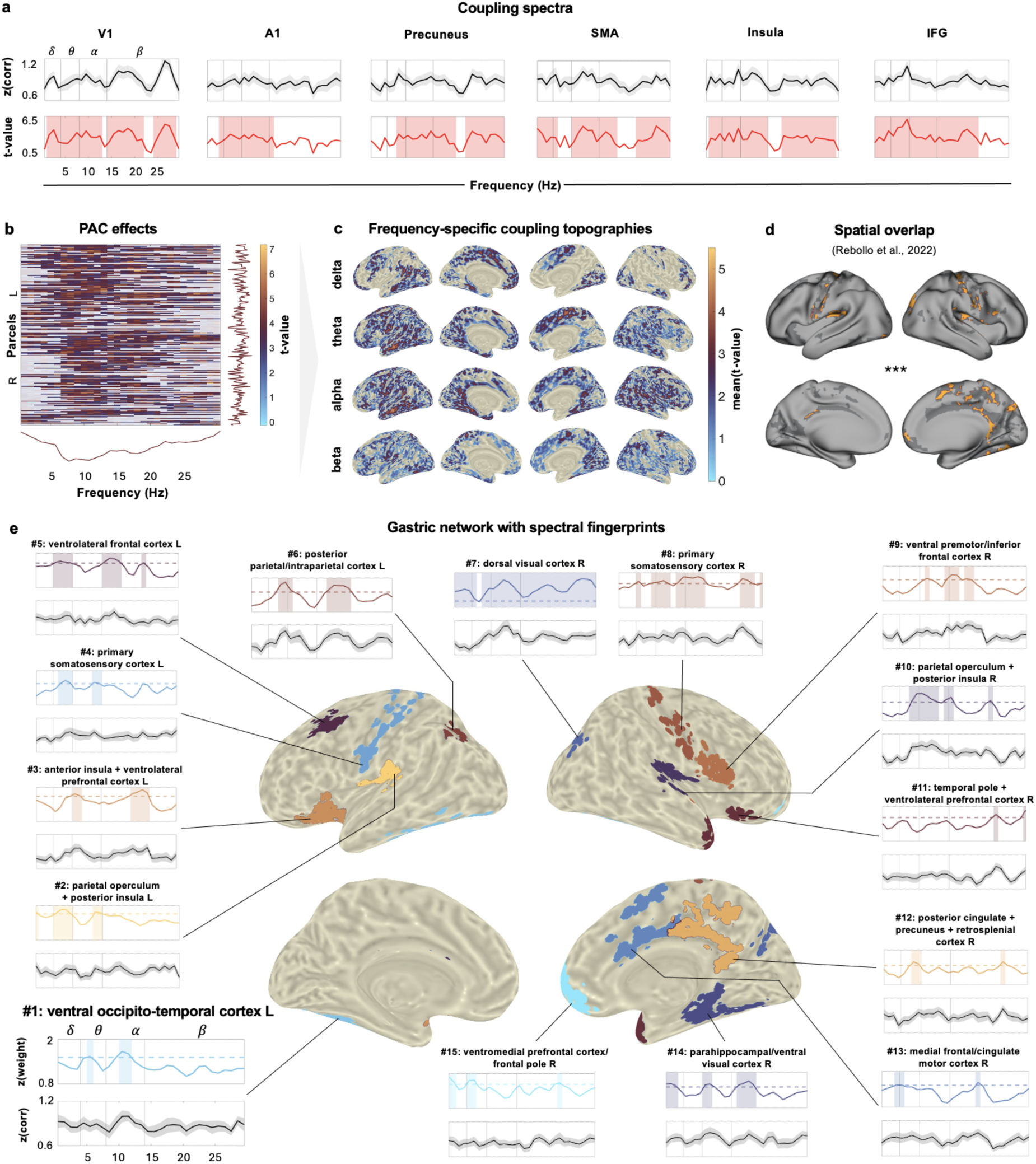
Gastric-brain coupling at rest. **a**, Fisher-transformed correlation and t-value coupling spectra for exemplary parcels in the right hemisphere, frequency ranges with significant coupling determined in cluster-permutation test indicated by red shading. **b**, Results for whole-brain cluster permutation test of correlation coupling spectra, non-significant frequencies & parcels opaque. Density curves depict the sum of significant t-values across parcels and frequencies, respectively. **c**, Frequency-band specific coupling maps. Source-level coupling projected to surface, significant t-values averaged within each frequency band. **d**, Significant spatial overlap of gastric network identified in this study using spectral analyses vs. gastric resting state network based on N = 63 fMRI participants described by Rebollo and Tallon-Baudry^12^. Overlapping regions marked in orange, dark grey denotes areas exclusively found in one of the two network maps. **e**, Gastric network components identified using NMF with their spectral fingerprints. Upper panel shows z-scored, mean component weight spectra. Frequency ranges showing consistent strong expression across subjects in the component are shaded. Dotted line marks permutation-based 95th percentile threshold of chance-level weights. Lower panel depicts the mean component Fisher-transformed correlation spectra, SEM shading across subjects.

In line with previous MEG work by Richter and colleagues^13^, we found the strongest coupling in the alpha frequency range (8 – 13 Hz). Core foci of gastric phase-modulated alpha power were the left temporo-insular region, the right V3, the left temporo-parietal-occipital junction, the left parahippocampal cortex, and the right medial premotor cortex, with alpha modulation showing the largest spatial spread, encompassing nearly all cortical regions. However, canonical delta, theta, and beta frequency ranges also showed robust modulation across both unimodal and transmodal areas (cf. Fig. 2b and c). In the delta band (0.5 – 4 Hz), we observed the strongest modulation in the left temporo-parietal-occipital junction, right V4, left frontopolar prefrontal cortex, left inferior parietal cortex, and left mid-cingulate cortex. In the theta band (4 – 8 Hz), peak modulation occurred in the left temporo-insular region, right V3, left superior parietal cortex, left superior temporal sulcus, and left secondary somatosensory cortex. Within the beta range (13 – 30 Hz), we found the strongest coupling effects in the left superior parietal cortex, right medial premotor cortex (SMA), left superior parietal somatosensory association cortex, right second visual area, and again the left superior parietal cortex. As illustrated by both the effect sum curve and frequency-specific coupling effect maps, gastric-brain coupling did not follow a clear pattern across the cortical hierarchy (Fig. 2b and c).

To ensure the specificity of the observed effects to the gastric rhythm, we repeated the PAC analysis after filtering each subject’s EGG signal at frequencies slightly lower and higher than the individual peak frequency (± 0.003 Hz, respectively). Coupling effects were significantly reduced for these filter-shifted control signals (*p* < .001 for both positive and negative shift vs. original frequency, see Supplementary Fig. 1a), confirming that the broadband modulation we observed was indeed specific to the normogastric frequency of the stomach. Because the PAC metric applied here is based on power values, we additionally tested whether the observed modulation could be explained by baseline power differences between parcels. Parcel-wise PAC effect sizes did not correlate with the respective baseline power, indicating that the observed effects were not driven by power-related biases (all *p* > .05, FDR corrected; Supplementary Fig. 1b).

### Spectral fingerprints of the gastric network

After establishing the spatial and spectral scope of gastric modulation of spontaneous oscillatory activity across the brain, we next aimed to compare our results to the gastric network previously identified in resting-state fMRI studies^11,12^. To this end, we extracted a spatially constrained map of components capturing the strongest coupling effects by applying non-negative matrix factorization (NMF^18^) to the parcel-wise PAC correlation spectra (0.5–30 Hz; see Methods). This allowed us to characterise fingerprints of frequency-specific gastric modulations of brain oscillations across the cortex.

Using this approach, we obtained a low-dimensional representation comprising 15 components per participant, parcel, and frequency, reflecting gastric modulation of oscillatory amplitude. Each component’s spatial map was thresholded at the 99th percentile, isolating the parcels exhibiting the strongest modulation for each component. In the following, we refer to the resulting anatomical map of components as a ‘network’; however, it should be noted that this is not equivalent to a network derived from classical connectivity analyses. We then quantified the correspondence between our oscillatory-derived network to the spatial distribution of the gastric resting state network described by Rebollo and Tallon-Baudry^12^. As shown in Fig. 2d, the map of components identified here showed large significant spatial overlap with the gastric resting state network (Dice similarity = 0.236, *p* = .001; spatial rotation permutation test), highlighting the robustness of gastric-brain coupling across different methodologies and timescales.

In addition to replicating the spatial organisation of the gastric network, we then leveraged the dimension-reduced NMF representation of broadband coupling effects across the cortex to characterise each component’s spectral profiles. We performed a non-parametric permutation test to quantify the frequency-specific consistency of component modulation across subjects. Thresholding the component weight spectra at the 95th percentile of chance-level consistency yielded a “spectral fingerprint” for each component. Consistent frequency ranges differed across components, indicating distinct, region-specific patterns of gastric-modulated oscillations. Taken together, the observed effects were relatively evenly distributed across theta, alpha, and beta frequency bands, with fewer components showing strong modulation in the delta range.

The anatomical location, mean coupling correlation spectrum, and frequency profile of each component are depicted in Fig. 2e. Overall, the cortical map of gastric phase-modulated oscillations prominently featured nodes of previously established brain-body networks^1^, including the insula, posterior cingulate cortex, and ventromedial prefrontal cortex. The insula (components #2-3 and #10) showed consistent modulation primarily in the theta, alpha, and beta bands. The posterior cingulate cortex (#12) exhibited above-chance modulation predominantly in the beta range (20 – 25 Hz). The ventromedial prefrontal cortex (#15) showed consistent modulation in both delta and beta bands. Notably, the dorsal visual cortex (#7) demonstrated the strongest effect, with above-chance consistency across the entire frequency spectrum.

### Global preferred phases of gut-related spectral changes

So far, we have identified a network of gastric-modulated brain oscillations using circular-linear correlation and dimensionality reduction. Given the distinct region-specific frequency profiles observed above, we next assessed whether the underlying modulation mechanism is shared across the brain or varies as a function of brain region. Our approach using phase-resolved power spectra to quantify PAC allowed for the analysis of power fluctuations across the gastric cycle, as well as determining the preferred phase with the maximal modulation effect.

As illustrated by the phase-resolved spectra and power changes across phase bins in Fig. 3a, oscillatory power across the gastric cycle followed a similar pattern across the whole brain, with the maximal power occurring in the upstate phase and the minimal power during the downstate phase of the gastric rhythm. A Rayleigh test on the circular means across phase bins identified the transition of falling to rising phase as the globally preferred phase of modulation (circular mean = – 3.12, *p* < .001, *z* = 229.65, see Fig. 3b).

**Fig 3.**
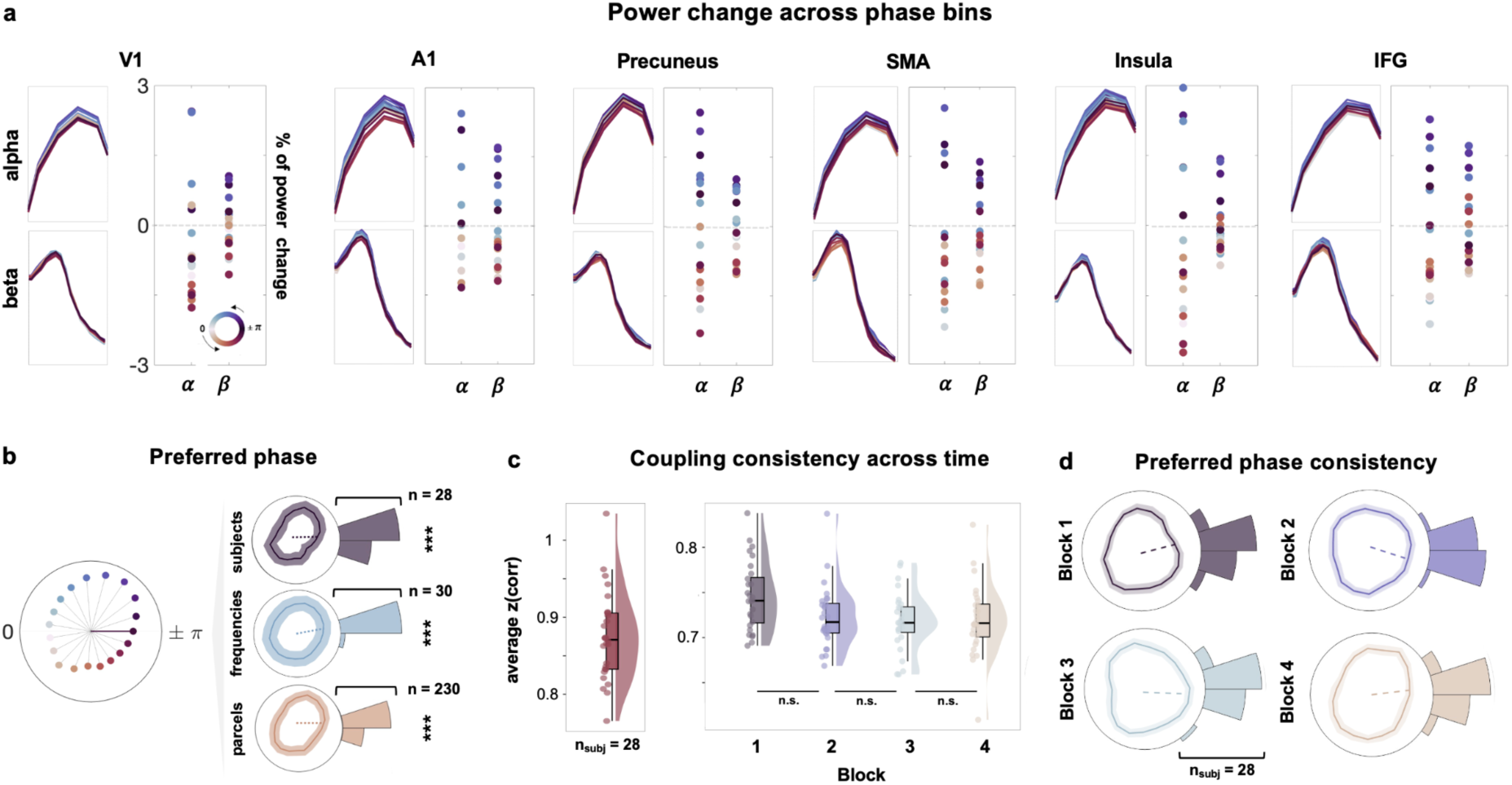
Modulation pattern consistency. **a**, Power changes across phase bins. Phase-resolved power spectra for alpha and beta frequency bands in same exemplary parcels from figure 2 panel a, averaged across hemispheres. Scatter plots illustrate percentage change of power across gastric cycle, zero denotes mean amplitude across all gastric phase bins. **b,** Preferred phase analysis. Polar plots display mean power across the gastric cycle with SEM shading; circular mean denoted by dotted line. Outer polar histograms show distribution of preferred phase values. Asterisks mark significant phase-locking across subjects, frequencies, and parcels, respectively. **c,** Coupling consistency across time. Left raincloud plot shows subject-level mean coupling strength across all frequencies and parcels, computed over the entire recording. Right panel illustrates consistency of subject-level mean coupling effects across four temporal quartiles. **d,** Consistency of preferred phase across time. Mean preferred phase across frequencies and parcels calculated for each subject for temporal quartiles. Polar plots show mean power across the gastric cycle with SEM shading. Circular mean denoted by dotted line, histograms denote preferred phase value distributions across subjects. SMA = supplementary motor area, IFG = inferior frontal gyrus.

To further evaluate the robustness of this phase preference, we also performed Rayleigh tests across frequencies and subjects, respectively. Both analyses revealed the same common preferred phase across brain regions, frequencies, and subjects (subject circular mean = – 3.12, *p* < .001, *z* = 27.68; frequency circular mean = – 3.12, *p* < .001, *z* = 29.98, see Fig. 3b).

These results point to a shared underlying mechanism in which gastric–brain coupling is preferentially associated with the transition between the end of a stomach cycle and the beginning of the next one. However, the phase of the gastric rhythm is partly dependent on the specific anatomical recording location: gastric contractions propagate as a travelling wave along the stomach, resulting in small spatial phase shifts (cf. supplementary Fig. 1c). To ensure the observed effects were not driven by such location-dependent, individual shifts in the analysed gastric phase signals, we computed the maximum observed phase shift between the most distant electrodes in the high-density EGG array. The mean maximum phase shift was 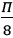, corresponding to a maximum shift of one phase bin, which is too small in magnitude to introduce meaningful variance in the preferred phase analyses. Moreover, the electrode selected for analysis was spatially consistent, with substantial overlap across subjects (see heatmap in supplementary Fig. 1d).

### Gastric-brain coupling is consistent across time

Having established the spatio-spectral organisation of gastric-modulated brain oscillations and a consistent underlying modulation mechanism, we lastly examined their temporal dynamics. Any coupling measure – both the PAC measure applied here as well as all previous measures used to quantify gastric-brain coupling^11,12,13^ – is computed on data aggregated across the entire recording duration. Consequently, the observed effects do not permit inferences about potential temporal fluctuations in coupling strength, including the possibility that effects are driven by brief epochs of particularly strong coupling.

To assess the temporal stability of gastric phase–dependent modulation of oscillatory amplitude, we segmented the recordings into overlapping windows comprising approx. 10 gastric cycles and recomputed PAC strength and preferred phase for each segment. Coupling strength was consistent across time (multivariate dependent-samples ANOVA, all *p* > .05, cluster-corrected, Fig. 3c). Similarly, preferred phase did not differ significantly across the segments (Watson-Williams test, *F*(3, 108) = 0.49, *p* = .69, Fig. 3d). These results indicate that gastric-brain coupling is a temporally sustained phenomenon rather than a sparse event.

### The gastric rhythm shapes directed network connectivity

In sum, our results thus far indicate that broadband neural oscillations are systematically coupled to the gastric rhythm through a consistent, temporally stable underlying mechanism. In a final step, we aimed to elucidate whether the gastric rhythm modulates functional connectivity within the previously identified component network using pairwise multivariate Granger causality (mGC). This approach allowed us to characterize gastric modulation of frequency-specific changes in communication between brain regions in the gastric network. Following the same logic as for the phase-resolved power spectra, we estimated time-resolved, pairwise mGC spectra in the frequency range between 0.5 and 30 Hz for all network component pairs using the two parcel time series within each network component and related them to gastric phase (see Methods for details). We then conducted a first-level linear regression predicting GC per frequency and pair of components from gastric phase, and computed the directed asymmetry index (DAI) for the resulting regression weights as a measure for frequency-specific directionality between any two components. As positive DAI values here indicate a stronger gastric modulation of *A* → *B* connectivity and negative values indicate a stronger gastric modulation of *B* → *A* connectivity, we tested the gastric regression weights DAI against zero using a two-sided test with cluster-based permutation correction. Note that this test assesses the difference in gastric modulation between the two directed connections; thus, a nonsignificant result establishes neither that gastric phase has no effect on either connection nor that modulation is equal.

On the group level, this revealed gastric phase-modulated broadband directed connectivity within the whole component network, with each component showing gastric-modulated information flow asymmetry for at least one connection (Fig 4b, right panel). Figure 4c illustrates the changes in absolute DAI values across gastric phase bins for these significantly modulated connections, separately for each frequency band. Analysis of the preferred phase across frequencies and pairs (Fig. 4c, right panel) identified a consistent pattern of connectivity strength and directionality across the gastric cycle highly similar to the oscillatory power changes presented above (Rayleigh test, frequencies circular mean = –3.11, *p* < .001, *z* = 6.85, pairs circular mean = –2.92, *p* < .001, *z* = 58.92). Across all frequencies and component pairs, the maximal directional influence occurred in the upstate phase and the minimal directional influence in the downstate phase of the gastric rhythm, further supporting an underlying mechanism of gastric-brain interaction preferentially associated with the transition between two gastric cycles.

**Fig 4.**
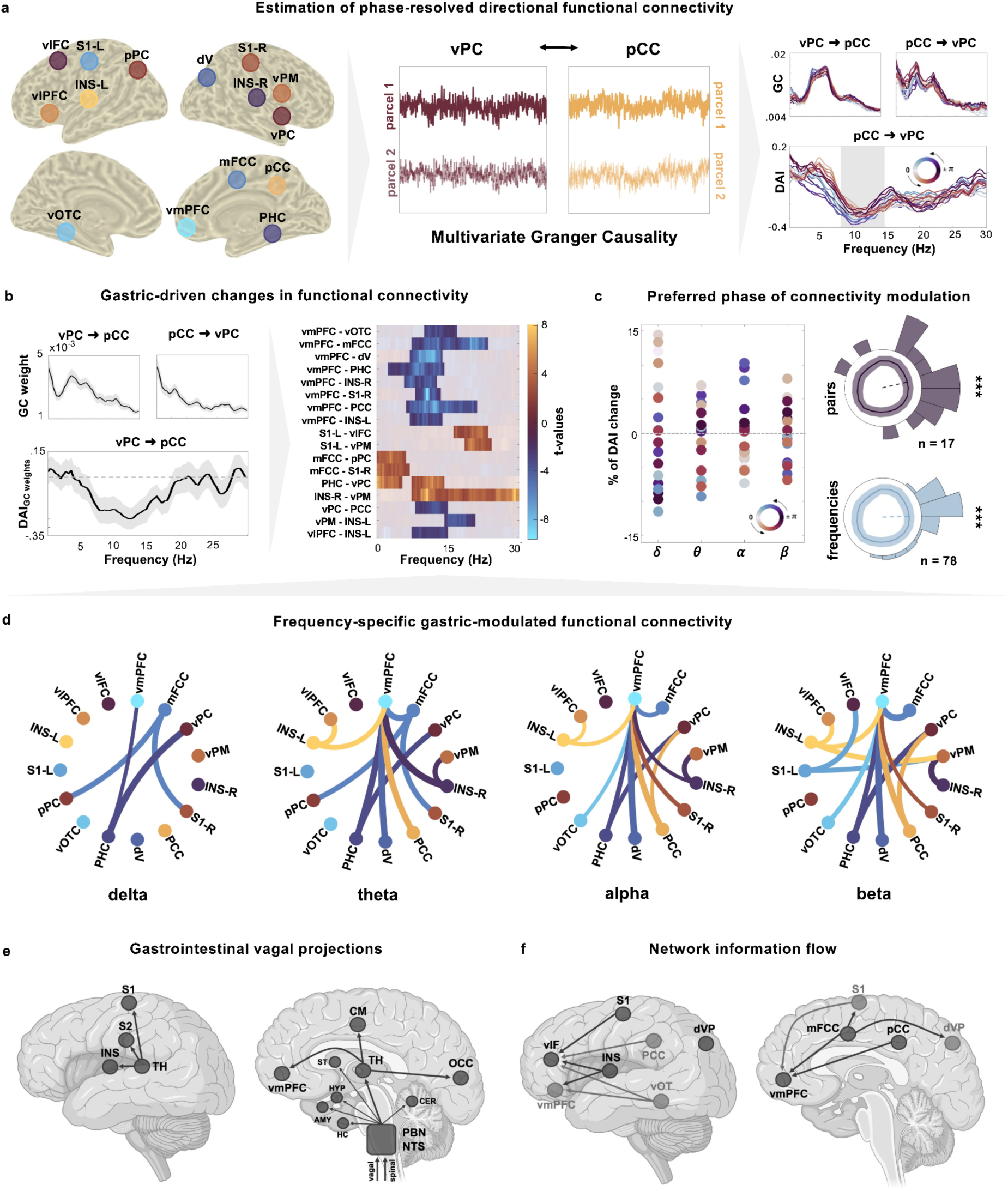
Gastric-modulated functional connectivity. **a**, Estimation of phase-resolved directed functional connectivity across the network components identified in the PAC analysis. Multivariate Granger (mGC) causality estimation and results for the exemplary network component pair ventrolateral prefrontal cortex R – posterior cingulate cortex R. The right panel shows gastric phase-resolved mGC spectra for both directions of information flow, as well as the phase-resolved DAI spectrum. **b**, Exemplary grand-average regression weight spectra between vPC and PCC in upper left panels (shaded areas show SEM). Lower panel shows regression weight spectra for vPC → PCC converted to DAI spectra (shaded areas show SEM). Right panel marks significant broadband gastric modulation of directed functional connectivity for pairwise conditions between network components. **c**, Percentage DAI changes across gastric phase bins for significantly modulated connections. Polar plots on the right show phase-resolved DAI across the gastric cycle, with polar histogram denoting the preferred phases of modulation across network component pairs and frequencies. Dotted line shows grand-average circular mean. Asterisks indicate significant clustering of preferred phases around a common preferred phase. **d**, Connectograms for gastric-modulated connections between network components in each frequency band, line color indicates an outgoing connection, and line width indicates modulation strength. **e**, Gastrointestinal vagal projections, adapted from Wolpert et al. (2020). **f**, Schematic of gastric-modulated network information flow as determined by gastric mGC modulation in this study.

A closer look at the frequency-band specific connections revealed a spatially and directionally organized frequency gradient (see Fig. 4d). In the delta and theta frequency bands, the gastric rhythm primarily modulated the directed influence from medial frontal and cingulate components toward posterior parietal and primary somatosensory regions. In the alpha and beta bands, gastric-modulated connectivity converged on the vmPFC, which received directed input from sources across the whole cortex, including pCC, PHC, mFCC, vOTC, INS, dorsal visual regions, and S1. A second, distinct alpha/beta cluster comprised somatosensory sources (posterior INS, S1) targeting ventrolateral and premotor frontal regions and the anterior insula. Together, these connectivity patterns suggest that gastric phase modulates distinct frequency-specific communication pathways: Slower rhythms primarily organize interactions along a cingulo-opercular axis, whereas alpha– and beta-band activity may coordinate the flow of somatomotor– and visual-system input toward higher-order associative regions, including the vmPFC.

Lastly, to place our findings of gastric-modulated cortical information flow in a physiological context, Fig. 4e and 4f depict the cortical targets of the ascending vagal gastric pathway (adapted from Wolpert *et al.*, 2020^16^) alongside a schematic of the gastric-modulated cortical connectivity patterns identified in this study. Vagal projection targets include the insula, S1, S2, cingulate motor areas, the occipital cortex, and the vmPFC. Notably, several of the projection targets – including the posterior insula, S1, cingulate motor cortex, and occipital cortex – appeared as source regions of gastric-modulated information flow in our results. The vmPFC however, although also a projection target, was observed as the primary convergence hub in our analyses. While this qualitative comparison does not establish a direct anatomical or physiological link, the observed correspondence is consistent with a possible physiological basis for the gastric modulation of directed information flow within the gastric network, with cortical sites receiving ascending gastric input acting as the source of gastric modulation of cortico-cortical communication.

## Discussion

In the present study, we provide the first comprehensive spatiotemporal mapping of human electrophysiological gastric-brain coupling at rest. Using a novel analysis approach based on circular-linear correlations of gastric phase-resolved power spectra, we report broadband coupling of the gastric rhythm with source-level neural oscillations across the cortex. Dimensionality reduction revealed distinct spatiospectral modulation patterns across the brain, mapping precise regional and spectral fingerprints of stomach-brain synchrony that highly overlapped with the gastric resting state network derived from fMRI by Rebollo and Tallon-Baudry^12^. Leveraging the advantage of high-resolution MEG recordings, we observed that the modulation patterns were not only highly consistent over time, but clearly showed a preferred gastric phase consistent across brain regions, frequencies, and participants. In the following, we will integrate our novel results with existing literature, highlight novel methodological developments, and propose a framework for interpretation.

Based on both animal and human studies, gut-brain communication is currently assumed to be mediated by vagal pathways transmitting information from the stomach to the cortex^19,20^. The corresponding cortical networks are largely based on fMRI recordings which provide high spatial resolution at the cost of temporal dynamics of stomach-brain coupling, which have so far been neglected. Here, the temporal precision of MEG allowed us to identify broadband gastric modulation of brain oscillations across both unimodal and transmodal regions including insula, primary and secondary somatosensory cortices, cingulate motor regions, frontal and prefrontal regions, and the ventral and dorsal occipital cortex. These regions not only span multiple levels of cortical organisation from primary sensory to higher-order integrative areas, but also precisely reflect cortical targets of vagal projection via the nucleus of the solitary tract, parabrachial nucleus, and the thalamus^1,16,21,22^.

In addition to receiving vagal afferents, they share a role in the processing and integration of visceral signals. The insular cortex serves as a central hub for interoceptive processing^23^, cingulate motor regions integrate visceral input with motor control^24^, and the ventromedial prefrontal cortex (vmPFC) represents a key hub of the default mode network coordinating visceral responses^25,26^. The occipital cortex has shown activation during gastric distension^27–29^, while somatosensory cortices have been implicated in parasympathetic and sympathetic control of the stomach^30^. Beyond this overlap in anatomical targets and functional roles, the spectral sensitivity of MEG revealed another shared feature: these regions exhibit dominant resting-state oscillations within the frequency bands most strongly modulated in our data. In the insula, beta and theta rhythms, thought to support intra-insular communication^31^, were preferentially modulated. In the vmPFC and cingulate cortex, modulation was most prevalent in the delta and theta bands, in line with their role as default mode network hubs and the established link between delta activity and default-mode network fluctuations^25^. Somatosensory cortices showed strong theta and alpha modulation patterns –frequency bands previously connected to the integration of multisensory information and bodily perception, respectively^32,33^. Finally, the occipital cortex, classically associated with alpha rhythm generation at rest^34^, showed broadband effects, suggesting that gastric modulation extends beyond canonical interoceptive networks.

Taken together, these observations suggest a fundamental role for stomach-brain coupling in cortical signaling at precisely the frequencies which determine region-specific transmission. This frequency-specific organization extends from regional oscillatory activity to interregional transmission, with gastric phase differentially shaping directed communication among coupled regions rather than exerting a uniform influence on functional connectivity. The modulated connectivity patterns suggest an influence of gastric phase on distinct communication circuits intersecting with canonical attentional, somatomotor, default-mode, and salience networks^12,35^. Consistent with the relationship between delta activity and default mode-network fluctuations^36^, gastric modulation of slower-frequency communication from medial frontal and cingulate regions to posterior parietal and primary somatosensory areas may support information flow along an attentional-somatomotor axis^37,38^. In contrast, the observed alpha– and beta-band modulation suggests an influence on the coordination of distributed salience-, somatomotor-, and visual-system inputs with higher-order associative regions, including the vmPFC, while also shaping communication within a second salience circuit^39–41^. Although it remains for future work to establish these network-level correspondences and their perceptual and behavioral consequences directly, their confirmation could extend recent evidence that bodily rhythms contribute meaningfully to perceptual processing by coordinating communication among interoceptive, sensory, attentional, and motor systems^42^. Considering the widespread beta-band effects, it is noteworthy that resting-state networks exhibit amplitude correlations most prominently in the beta frequency band^25^, suggesting that gastric phase may modulate interactions among intrinsic cortical networks. More broadly, the observed frequency gradient is compatible with proposed spectral asymmetries in cortical communication, whereby different frequency bands support distinct modes of interregional interaction^43–46^. Within this framework, gastric phase may contribute to the temporal coordination of intrinsic cortical processing and interregional communication at rest, with the alpha– and beta-band effects being consistent with proposed roles of these rhythms in information gating, top-down coordination, and the maintenance of ongoing network states^47,48^.

Anatomically, the spatial distribution of the observed directed connectivity effects is also notable: several sources of gastric-modulated information flow – including the posterior insula/operculum, S1, and cingulate motor cortex – have been identified as cortical targets of ascending vagal pathways^1,16,21,22,49^. While we discuss the question of causation in detail below, these findings tightly connect vagal afferents from the stomach to the brain and oscillatory dynamics supporting the cortical function at rest. Hence, the observed gastric modulation conceivably reflects the processing of visceral input from the stomach into the cortex, e.g. for the purpose of homeostatic and allostatic regulation. Aside from these effects on local cortical processing, the fact that broadband coupling effects extended across all cortical regions points to a role of gastric modulation beyond visceral processing alone.

Converging fMRI and MEG evidence demonstrates the robustness of stomach-brain synchrony effects across recording methods and timescales. Our spatially constrained map of dominant oscillatory coupling patterns across the brain and frequency spectrum overlapped significantly with the gastric resting state network comprised of brain regions whose fMRI signals are phase synchronized to the gastric rhythm^12^. Consistent with this overlap, the close correspondence between gastric-modulated frontal, cingulate, and parietal regions and canonical resting-state networks replicates Rebollo and Tallon-Baudry’s^12^ report of the gastric network cutting across multiple classical resting-state network nodes. Notably, resting-state network nodes exhibit amplitude correlations predominantly in the beta frequency band^36^, which was among the most strongly and consistently modulated frequency bands in our data. In line with earlier interpretations in the respiratory domain^5^, beta-band gut-brain coupling strongly suggests that dynamic interactions between resting-state networks are at least in part explained by physiological rhythms from the lungs and the stomach.

Interestingly, the congruence of gastric effects does not extend to the previously reported underrepresentation of modulation effects in transmodal areas^12^. As transmodal areas are central to large-scale integration and support inter-areal communication through oscillatory coupling across distributed brain regions^50–52^, the broadband gastric coupling effects observed here are consistent with a contribution of the gastric rhythm to large-scale oscillatory brain dynamics. Most likely, it requires the temporal precision of MEG recordings to capture the dynamics of these coupling effects over time, meaning they may only be partly reflected in the slower haemodynamic fluctuations measured by fMRI^53–55^.

As for the frequency specificity of these coupling effects, we also replicated the gastric-brain alpha rhythm modulation reported by Richter *et al.*^13^ in the anterior cluster including the anterior insula, inferior frontal gyrus, putamen, and superior temporal gyrus, as well as the posterior cluster including the parieto-occipital sulcus and the calcarine fissure. Despite using an exploratory, data-driven approach across brain regions and frequencies, Richter *et al.*’s^13^ results were limited to a rather narrow alpha frequency range of 10-11 Hz. The disparity with our broadband, widespread effects can be explained by differing methodological approaches: Richter *et al.*^13^ quantified coupling between the gastric rhythm and neural oscillations using the Modulation Index^56^ and statistically tested against a chance-level distribution based on time-shifted surrogate gastric signals. However, circular shifting of phase information does not entirely destroy phase-amplitude coupling in the null distribution, which compromises statistical inference. To mitigate this limitation, we applied a novel approach using circular-linear correlations of gastric phase and neural power. Our reasoning was that shuffling phase labels on the phase-resolved spectra allowed us to truly disrupt amplitude-phase relationships, resulting in a true chance-level distribution for statistical testing.

Combined with the high temporal precision of MEG recordings, this new approach allowed us to characterise a global modulation pattern of neural oscillations across the gastric cycle. The gastric phase associated with maximal power and the preferred phase of PAC was congruent across subjects, brain regions, and time; with the same phase preference also extending to directed interregional communication, indicating one common underlying mechanism of gastric modulation of brain oscillations and within-network communication. At the neurochemical level, all neuromodulatory systems could be considered plausible candidate mechanisms – gastric afferents reach dopaminergic, noradrenergic, histaminic, cholinergic and glutamatergic systems^20^. Notably, activity in the locus coeruleus – the principal source of forebrain norepinephrine – is modulated by gastric distension^57^. At the physiological level, the efferent vagal projections from the stomach with their widespread cortical and subcortical targets represent a likely pathway for this mechanism. In rats, vagotomy (surgical disconnection of the vagus nerve) significantly reduces the coherence between brain activity and gastric slow waves^19^. In humans, recent evidence shows that vagal nerve stimulation can alter gastric rhythm frequency and amplitude^58,59^, and that gastric electrical stimulation enhances vagal activity^60^, presenting promising interventions to establish causal direction and elucidate the physiological mechanisms of stomach-brain interactions in humans.

Despite the robustness of the spatial distribution of gastric-brain modulation across recording modalities, our findings of a global preferred gastric phase across the cortex seem to contradict the gradient phase shift reported in previous fMRI literature at the first glance^11^. Yet, again considering the cross-methods differences in temporal resolution, this may simply reflect different timescales of the same process (fast PAC dynamics captured by MEG vs. slower metabolic activity in BOLD signal), emphasising the need to investigate stomach-brain interactions across different timescales and neuroimaging methods.

Furthermore, our results show a specific preferential association of the modulation with the transition between the end of one and the beginning of the next stomach wave. In this context, it is noteworthy that the gastric phase is dependent on the anatomical recording location. Gastric slow waves propagate as a travelling wave along the stomach^61^, introducing a consistent phase shift across the array of recording electrodes. Thus, the exact gastric phase driving stomach-brain synchrony is location-dependent. Our application of a high-density EGG recording layout allowed us to disentangle true variance in preferred phase and variance due to recording location. The high-density layout not only improves the signal-to-noise ratio – by better accounting for individual differences in stomach anatomy when placing EGG electrodes based on anatomical landmarks and reducing participant dropout due to unusable signals – but also provides insights into the propagation of the gastric rhythm itself^62^. In our analyses, this allowed an estimate of the maximum possible variance in gastric phase introduced due to recording location by tracking the phase shift in the gastric time series across the electrode array. Nonetheless, we caution against interpreting the specific phase identified here as carrying physiological information or having a unique functional meaning. Rather, the significance of this finding lies in the temporal consistency of the phase preference. This distinction is captured by the two modes of brain-body interaction proposed by Tallon-Baudry *et al.* (2026)^63^: In the so-called ‘information mode’, bodily signals – and potentially specific timepoints within those signals – convey information about the organism’s internal state to the central nervous system. In ‘coordination mode’, by contrast, the rhythmic structure of these signals provides a temporal framework for organizing neural activity. We interpret our preferred-phase findings in the latter context: Rather than assigning specific informational content to the identified gastric phase, we suggest that the temporal structure of the gastric rhythm (captured by its phase) may provide a recurring temporal reference for coordinating distributed cortical dynamics.

Overall, cross-frequency coupling is widely regarded as a principle of temporal organization of global brain activity and between-area communication^64^. Through oscillatory synchrony, windows of more or less excitability are imposed, coordinating activity between distant groups of neurons^1^. In this work, we show that the temporal structure of broadband neural activity is modulated by the gastric rhythm. Within the framework of neural communication through oscillatory coherence, this suggests that the gastric-modulated areas are synchronized, with the gastric rhythm acting as a carrier wave. Given the strong and spatially widespread gastric modulation observed across the brain, we therefore propose that the gastric rhythm may contribute to large-scale brain organisation by providing a scaffold for oscillatory synchrony^1,64,65^. The phase-dependent modulation of directed information flow further indicates that this scaffold may extend to the functional organization of interregional communication, with the gastric rhythm serving as a temporal reference signal that coordinates the availability of cortical communication channels. Consistent with this scaffolding hypothesis, we observed strong phase coherence of gastric modulation across brain regions, frequencies, and participants, indicating a shared global mechanism. Moreover, the observed coupling effects were highly stable over time, further supporting the notion that gastric modulation reflects a persistent organisational feature. The evolutionary conservation of the frequency of both intrinsic brain rhythms and the gastric rhythm^16,66^ suggests that this mechanism may even translate across species. For example, Cao *et al.*^19^ recently identified a network of stomach-brain phase coherence in the rat brain, and Rafael *et al.*^67^ demonstrated a neuronal basis for the perception of internal gut sensations in the rat insular cortex.

However, it remains unclear whether this bodily scaffold has direct functional consequences or primarily serves as a ‘metronomic’ signal that facilitates neuronal synchronization by imposing rhythmic windows of excitability. Recent findings link increased frontoparietal gastric-brain coupling to poorer mental health^68^, and the gastric rhythm modulates reaction times across different behavioral paradigms^69^, providing initial evidence for a gastric influence on behavior. Further, altered gastric interoception has been implicated in eating disorders^70^. Ultimately, further systematic investigations of gastric-behavioral interactions, as well as gastric modulation of neural activity across healthy and clinical populations and varying brain and body states are required to establish the functional consequences of gastric scaffolding of brain dynamics.

Lastly, the gastric rhythm is but one among many visceral rhythms present in the body. It remains an open question how gastric-brain scaffolding integrates with other bodily rhythms also known to shape the temporal organisation of brain dynamics^5,71^. Future work should therefore broaden the scope and adopt a higher-order model of body-brain interactions across multiple physiological rhythms, including cardiac and respiratory signals, to truly understand the impact of visceral signals on neural function and cognition^72^. Despite consistent reports of considerable anatomical overlap in the cortical targets of these rhythms^1,12,13^, understanding how they interact – whether they nest, reinforce, or compete to organize neural activity – remains one of the biggest challenges in the field today.

## Methods

### Participants

In total, 30 volunteers (15 female, *M_age_* = 25.5, *SD_age_* = 2.8 years) participated in this study. All participants had very good German language skills, denied having any neurological diseases, and gave written informed consent before participating. The study was approved by the ethics committee of the University of Münster (ethics ID 2019-713-f-S.).

### Procedure

For resting state recordings (duration 14min), participants were seated upright in a magnetically shielded room with their eyes open, fixating a central marker on the screen in front of them. MEG and EGG data were recorded simultaneously in a continuous recording at a sampling frequency of 600 Hz (EGG with DC recordings and no low-pass filter). MEG data was acquired using a 275-channel whole-head system (OMEGA 275, VSM Medtech, Vancouver, Canada). The participants’ heads were stabilised with cotton pads inside the MEG helmet to ensure minimal head movement.

To record the gastric rhythm, a high-density EGG with an array of 20 cutaneous, passive electrodes was placed on the participants’ abdomen. Reference and ground electrode of a simultaneous ECG recording were used as reference and ground electrode for the EGG electrodes, respectively (Fig. 1a). For easier access to anatomical landmarks, electrode placement on the participant’s abdomen was carried out sitting in a reclined position. First, a central point on the midline of the abdomen was determined by vertically moving downwards from the xiphoid process towards the umbilicus until the horizontal distance from midline to left and right ribcage measured 5 cm. Two electrodes were placed in 5 cm horizontal distance to the right and left of this point respectively, directly adjacent to the ribcage. These two electrodes formed the upper corners of the electrode array. Next, the point on the midline 2 cm above the umbilicus was identified and two more electrodes were added to the left and right with 5 cm horizontal distance, completing the lower corners of the electrode array. The space between the upper and lower corner electrodes was then filled to form an equidistant array of 4 rows x 5 electrodes.

For the purpose of MEG source localization, high-resolution structural magnetic resonance imaging (MRI) scans were acquired in a 3T Magnetom Prisma scanner (Siemens, Erlangen, Germany). Anatomical images were obtained using a standard Siemens 3D T1-weighted whole brain MPRAGE imaging sequence (1 × 1 × 1 mm voxel size, TR = 2130 ms, TE = 3.51 ms, 256 × 256 mm field of view, 192 sagittal slices). Scans were conducted in supine position to reduce head movements, and gadolinium markers were placed on the nasion as well as the left and right distal outer ear canal positions for landmark-based co-registration of MEG and MRI coordinate systems.

### MEG processing

All data was preprocessed using the Fieldtrip toolbox^73^ running in MATLAB 2024b (The MathWorks Inc., 2024). MEG data were demeaned, detrended, resampled to 300 Hz, and visually inspected for movement and technical artefacts. Segments of data marked as artefacts were rejected and removed. Subsequently, the time axis of the data was reconstructed to retain its original continuous structure. MEG data were then noise balanced and a discrete Fourier transform (DFT) filter was applied to remove 50 Hz line noise and its harmonics. A visual inspection based on channel amplitude variance and kurtosis identified bad channels which were then repaired using spline interpolation of neighbouring channels (*M*_rejchans_ = 3.73, *SD*_rejchans_ = 4.12). Lastly, independent component analysis (ICA) using the *runica* algorithm was performed to visually exclude eye and cardiac components, resulting in a range of 1 to 5 components being rejected per participant (*M*_rejcomps_ = 3.01, *SD*_rejcomps_ = 1.45).

### EGG preprocessing

EGG data were demeaned, detrended, and resampled to 300 Hz. A fast Fourier transform with a Hanning taper (to reduce spectral leakage and control frequency smoothing) was applied to identify both the channel with the peak power value as well as the corresponding peak frequency in the normogastric range between 0.033 and 0.067 Hz. Identifying one single channel with peak power to determine gastric frequency accounts for intersubject variability in precise anatomical stomach location and yields a better signal-to-noise ratio. The time course of the identified peak channel was filtered using a FIR2 filter with a bandwidth of 0.015 Hz around the identified peak frequency. Further, a Hilbert transform was applied to the filtered time course to extract instantaneous phase and amplitude of the EGG signal.

Visual inspection of power spectral density across all electrodes, filtered time course of the peak power electrode as well as the time course of the instantaneous phase ensured data quality in line with the recommendations on EGG data quality in Wolpert *et al.*^16^.

### MRI co-registration and extraction of time series in source space

Co-registration of structural MRIs to the MEG coordinate system was done individually by identification of three anatomical landmarks – nasion, left and right pre-auricular points – in the participant’s MRI. Individual head models were constructed from the anatomical MRIs using segmentation algorithms implemented in Fieldtrip and SPM12 (SPM Team, University College London, 2014). A solution of the forward model was computed using the realistically shaped single-shell volume conductor model^74^ with a 5-mm grid defined in the Montreal Neurological Institute (MNI) template brain (MNI, Montreal, Canada) after linear transformation to the individual MRI. Source reconstruction was performed using the linearly constrained minimum variance (LCMV) beamformer approach^75^ with the regularisation parameter set to 5%, estimating a three-dimensional spatial filter for each location of the 5-mm grid. Spatial filters were then combined into a reduced grid of three-dimensional filters corresponding to the 230 parcels of a recent multimodal parcellation of the cerebral cortex^17^ by using the covariance matrix of all filters in each parcel to perform a singular value decomposition. The filters for each parcel were then further reduced in dimension to the direction yielding maximum power. A single broadband LCMV beamformer in the range of 0 – 150 Hz was used to estimate the parcel-level time course across all frequencies.

### Computation of phase-amplitude coupling

We computed PAC as the circular-linear correlation of phase-resolved, frequency-specific neural power with the corresponding gastric phase angle and generated a null-distribution for statistical testing by shuffling phase labels of the phase-resolved power spectra.

In a first step, we removed frequency bias from the source parcel time series by considering the first-order derivative, which can be used as a time-domain filter of the aperiodic component^76^. Time-resolved power was obtained for 30 equally spaced frequencies in the range of 0.5 – 30 Hz using a moving FFT window approach (2-second windows with 50% overlap). The power spectra were then binned into 20 gastric phase bins, resulting in an average phase-resolved power spectrum for each parcel (Fig. 1b). Accounting for the circular nature of gastric phase data, we calculated single-subject level PAC for each parcel and frequency as the circular-linear correlation between gastric phase and binned power using the *circ_corrcl* function from the *circstat* toolbox for Matlab^77^. When the amplitude of a faster signal is modulated by the phase of a slower signal, power will vary systematically across the phase cycle, resulting in a high circular-linear correlation value (Fig. 1c). In the absence of PAC, average power values are randomly distributed across the phase cycle, leading to a low correlation value (Fig. 1c). This approach using circular-linear correlations has previously been successfully used by our group to quantify respiration-brain PAC dynamics^78^.

To determine the group-level significance of stomach-brain coupling effects, we created a null distribution of 500 chance-level correlation values by randomly shuffling the phase bin labels of the phase-resolved power spectra and re-computing circular-linear correlations on the shuffled data for each subject, parcel, and frequency. While this approach ensures that the original phase-amplitude relationship is truly disrupted for the chance-level null distribution values, shuffling labels on the level of mean phase-resolved spectra avoids the limitations of more conventional surrogate generation methods run on the time-series level (such as circular shifting or segment shuffling). Preserving the amplitude distributions is simple on both levels – however, the phase-resolved mean spectra no longer contain a temporal autocorrelation that needs to be respected, which allows for a completely random shuffling of phase without destroying inherent characteristics of the signal.

All empirical and chance-level correlation values were Fisher z-transformed prior to statistical testing. Significance of coupling effects was evaluated by comparing the 95th percentile of the null distribution values to the empirical correlation values for each subject using the non-parametric cluster permutation test procedure implemented in fieldtrip’s *ft_freqstatistics* dependent samples *t*-test with 5000 Monte-Carlo permutations (α = .05, right-tailed).

### Rank optimisation and NMF

One of the main aims of this work was the replication of the gastric network previously only identified in fMRI data using spectral analyses. Thus, we anatomically localized a network of gastric phase-dependent amplitude modulation effects using a spatially sparse variant of NMF. As the z-transformed correlation values are inherently positive, sparse NMF constituted an appropriate low-dimensional decomposition. Additionally, NMF was particularly well suited for our purposes of dimensionality reduction because it allows for an anatomically sparse mapping of components rather than preserving the full spatial topography – unlike methods such as k-means or hierarchical clustering. Another major advantage of the NMF approach is that it yields components with distinct, individual spectral profiles capturing the broadband PAC effects. This allowed us to characterise fingerprints of frequency-specific gastric modulations of brain oscillations across the cortex.

In more detail, we first smoothed the individual z-scored correlation spectra using a Gaussian kernel (window size 3.5 Hz) and then determined the number of components to be extracted with the NMF for each subject using Matlab’s *choosingR* function which evaluates optimum rank based on a singular value decomposition. Across subjects, this procedure yielded a median optimal number of 15 components. The smoothed correlation spectra were then reshaped to a group-level input matrix for the NMF (840 [30 frequencies x 28 subjects] x 230 parcels). On this matrix, we fitted the *sparsenmfnnls* from the NMF toolbox^18^ for Matlab with a sparsity parameter of 0.3, yielding the output matrices *A* and *Y*. The basis matrix *A* (840 [30 frequencies x 28 subjects] x 230 parcels) contains the subject-specific spectral profiles, quantifying each subject’s individual, relative contribution to each network component for each frequency. Coefficient matrix *Y* (15 components x 230 parcels) describes the spatial profile for each network component, representing each parcel’s relative contribution to each component. For a more detailed description of the sparse NMF algorithm refer to Li and Ngom^18^, a similar application of the algorithm in body-brain coupling can also be found in Chalas *et al.*^5^. While NMF results computed on a given data set will always be highly similar, any single solution depends on its random initialization. Therefore, we ran 100 iterations of the NMF analysis and selected the solution with the lowest residual error. Specific spatial locations for each component were obtained by thresholding the component contributions in the coefficient matrix *Y* at the 99th percentile, yielding a cortical map of NMF components corresponding to 30 atlas parcels in total. To assess how consistently frequencies were modulated on the group level within any given component, we used a non-parametric permutation procedure. After z-scoring the component weights across frequencies for each subject, we generated a null distribution of mean spectral values across all frequencies for each component by randomly shuffling the frequency axis of each subject’s component spectrum (5000 permutations). Thresholding the mean component spectra at the 95th percentile of this distribution yielded component-specific spectral fingerprints of gastric–brain coupling. Applying this procedure jointly across all components inherently provides multiple-comparison correction across both frequencies and components.

### Spatial overlap

To compare the spatial overlap between our findings and previous fMRI results on gastric-brain coupling^12^, we binarized the spatial map of network components derived from the NMF and transformed it to fsLR 32k surface space. Spatial overlap was then computed using the Dice similarity coefficient. We assessed statistical significance by generating a null distribution of Dice coefficients from 1000 spatially rotated surrogate maps^79^. P-values were calculated as the proportion of rotated Dice values exceeding the absolute empirical Dice coefficient.

### Modulation pattern and consistency analyses

We performed a series of analyses on the mean phase-resolved power spectra across subjects investigating the pattern of amplitude modulation across the gastric cycle. All functions related to circular statistics described in the following analyses were implemented using the *circstat* toolbox^77^. The percentage of source parcel power change for each gastric phase bin was calculated as the percent deviation from the mean amplitude across phase bins. Rank-order consistency of power across gastric phase bins was assessed using Friedman tests, computed separately across frequency bands within each parcel and across parcels within each frequency band. For this purpose, we averaged the phase-resolved power spectra within each frequency band and then normalised power across frequency bands to remove absolute power differences.

To identify the preferred phase of the gastric cycle driving the modulation, we calculated the circular mean of the phase-binned power spectra for each subject, parcel, and frequency using the *circ_mean* function. Rayleigh tests (*circ_rtest*) were used to determine whether the circular means clustered significantly around a specific phase, indicating consistent phase preference.

### Coupling consistency across time

To examine fluctuations of gastric-brain coupling across time, we divided the recordings into four chronological quartiles comprised of approx. 10 gastric cycles with 25% overlap and re-computed PAC effects as well as phase preference for each segment. We used a multivariate dependent-samples ANOVA with the cluster-based permutation approach implemented in fieldtrip’s *ft_freqstatistics* function to test for group-level differences in coupling strength across time blocks in each parcel and frequency (*ft_depsamplesFmultivariate*, 5000 Monte-Carlo randomizations, α = .05, right-tailed). Consistency of preferred phase angle across time was tested on subject-level block means with a Watson-Williams test using *circ_wwtest*^77^.

### Connectivity analyses

Multivariate Granger causality (mGC) between network components was computed using a pairwise, nonparametric approach^42,80^. First, we estimated the spectral representation of the source-localized time series from the two parcels comprising each component in the frequency range of 0 – 30 Hz, using the same moving-window multitaper Fourier approach described for the phase-amplitude coupling computation above. The resulting spectral estimates were again assigned to 20 gastric phase bins, and cross-spectral density matrices were computed separately for each component pair and phase bin. Based on these cross-spectral density matrices, we estimated mGC using a nonparametric spectral matrix factorisation in a blockwise approach, yielding phase-resolved spectra of directed information flow between any two network components. The multivariate computation allowed us to consider the combined influence of all parcel-level time series within each network component, providing a multivariate prediction of directed interactions rather than relying on a single parcel or component-averaged, unimodal signal.

To quantify gastric phase-locked modulation of directed functional connectivity in the gastric network, we then conducted a first-level linear regression predicting mGC per frequency and pair of components across gastric phase:

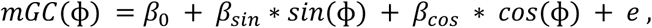

where ɸ denotes the gastric phase angle associated with each phase bin. The magnitude of phase modulation was then quantified by combining the regression weights in a gastric phase vector norm:

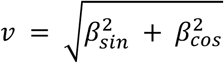

For each pair of components, we computed the directed asymmetry index (DAI) as a measure of frequency-specific directionality as follows:

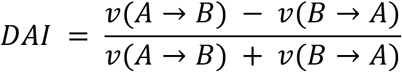

Positive DAI values indicate a stronger gastric phase modulation in the *A* → *B* direction, while negative values indicate stronger modulation in the *B* → *A* direction. Group-level significance of the DAI spectra was evaluated using the same cluster-based permutation test described for the phase-amplitude coupling spectra, with functional connectivity being considered modulated by gastric phase for frequencies and specific component pairs whose DAI was statistically significantly different from zero.

Lastly, phase-resolved DAI spectra for each component pair were calculated on the phase-resolved mGC spectra to assess the preferred phase of the modulation as well as its consistency across pairs and frequencies using the circular mean and Rayleigh tests, respectively.

## Acknowledgements

The authors thank Karin Wilken, Ute Trompeter, and Hildegard Deitermann for their essential assistance during data collection.

## Author contributions

Conceptualisation, D.S.K., I.R.; Methodology, D.S.K., T.B., Investigation, T.B., Writing – Original Draft, T.B., D.S.K., Writing – Review & Editing, T.B., D.S.K., I.R., M.S., J.F., Visualisation, T.B.; Funding Acquisition, D.S.K.

## Supplementary material

**Supplemental Fig S1.**
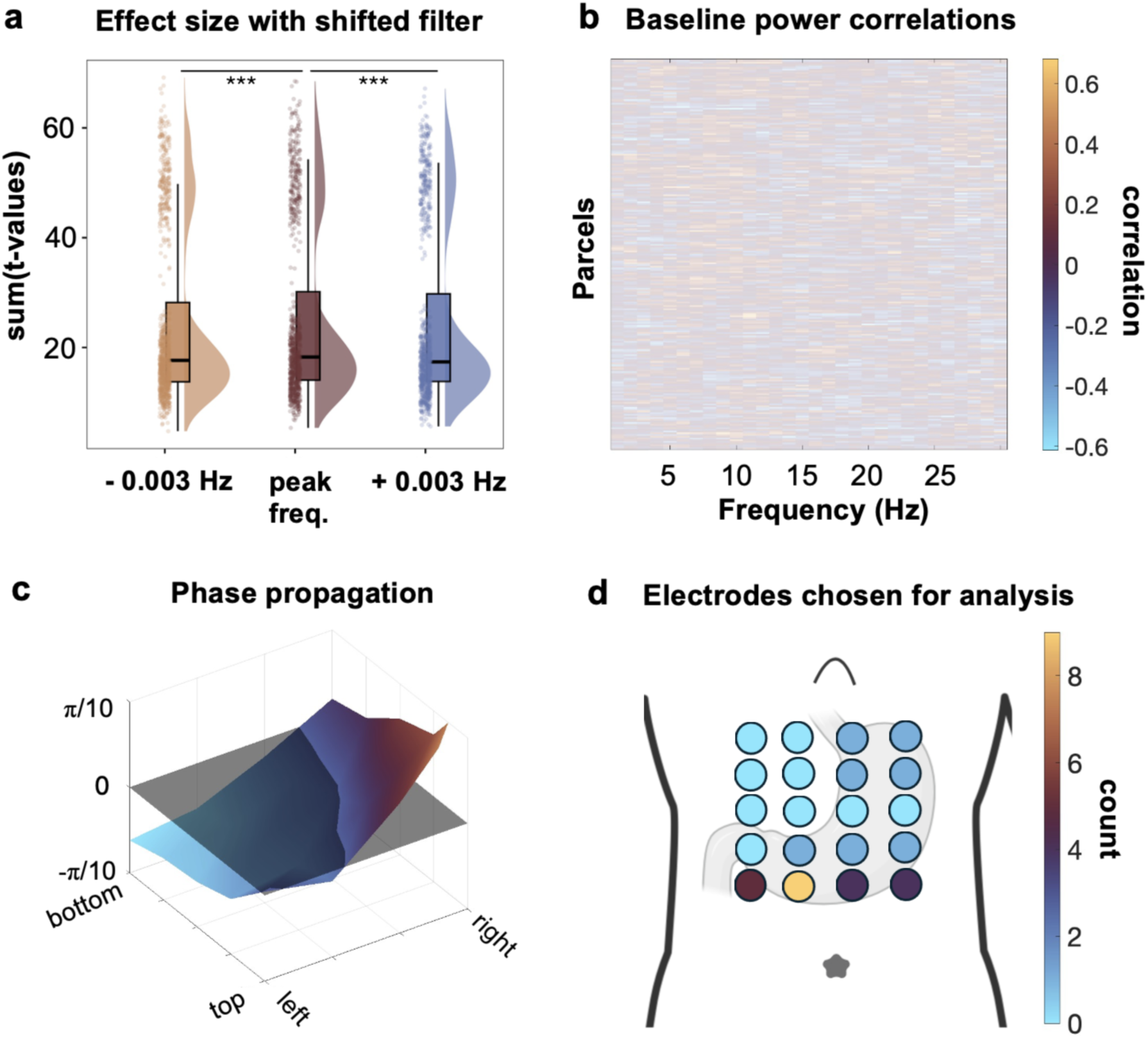
Control analyses. **a**, PAC effect sizes for shifted peak gastric rhythm frequency. PAC coupling analysis repeated after filtering EGG at centre frequencies slightly lower and higher than the individual peak frequency; shifts by 0.003 Hz in positive and negative direction, respectively. Summary t-values across subjects for each parcel and frequency depicted here. Significance of effect size differences between filter centres was assessed using a permutation test with randomly shuffled labels. **b,** PAC effects correlation with baseline power. Matrix shows correlation of PAC effects with baseline power across parcels and frequencies; significance corrected for multiple comparisons using FDR-BH. Non-significant parcels and frequencies opaque. **c,** Phase propagation along the stomach in high-density EGG layout. Exemplary data for one subject shown here. Grey plane denotes mean phase across all electrodes, coloured plane depicts phase deviations from mean for each electrode. Spatial layout corresponds to view of electrode locations in panel d. **d,** Schematic heatmap of electrodes chosen for analysis. Colormap indicates the number of times each electrode was selected for analysis across subjects.

## References

1. Engelen, T., Solcà, M. & Tallon-Baudry, C. Interoceptive rhythms in the brain. Nat. Neurosci. 26, 1670–1684 (2023).

2. Azzalini, D., Rebollo, I. & Tallon-Baudry, C. Visceral signals shape brain dynamics and cognition. Trends Cogn Sci (Regul Ed) 23, 488–509 (2019).

3. Tallon-Baudry, C. Interoception: Probing internal state is inherent to perception and cognition. Neuron 111, 1854–1857 (2023).

4. Parviainen, T., Lyyra, P. & Nokia, M. S. Cardiorespiratory rhythms, brain oscillatory activity and cognition: review of evidence and proposal for significance. Neurosci. Biobehav. Rev. 142, 104908 (2022).

5. Kluger, D. S. & Gross, J. Respiration modulates oscillatory neural network activity at rest. PLoS Biol. 19, e3001457 (2021).

6. Tort, A. B. L., Laplagne, D. A., Draguhn, A. & Gonzalez, J. Global coordination of brain activity by the breathing cycle. Nat. Rev. Neurosci. 26, 333–353 (2025).

7. Bozler, E. The action potentials of the stomach. American Journal of Physiology-Legacy Content 144, 693–700 (1945).

8. Suzuki, N., Prosser, C. L. & Dahms, V. Boundary cells between longitudinal and circular layers: essential for electrical slow waves in cat intestine. Am. J. Physiol. 250, G287–94 (1986).

9. Saper, C. B. The central autonomic nervous system: conscious visceral perception and autonomic pattern generation. Annu. Rev. Neurosci. 25, 433–469 (2002).

10. Rinaman, L. Hindbrain noradrenergic lesions attenuate anorexia and alter central cFos expression in rats after gastric viscerosensory stimulation. J. Neurosci. 23, 10084–10092 (2003).

11. Rebollo, I., Devauchelle, A.-D., Béranger, B. & Tallon-Baudry, C. Stomach-brain synchrony reveals a novel, delayed-connectivity resting-state network in humans. eLife 7, (2018).

12. Rebollo, I. & Tallon-Baudry, C. The Sensory and Motor Components of the Cortical Hierarchy Are Coupled to the Rhythm of the Stomach during Rest. J. Neurosci. 42, 2205–2220 (2022).

13. Richter, C. G., Babo-Rebelo, M., Schwartz, D. & Tallon-Baudry, C. Phase-amplitude coupling at the organism level: The amplitude of spontaneous alpha rhythm fluctuations varies with the phase of the infra-slow gastric basal rhythm. Neuroimage 146, 951–958 (2017).

14. Weaver, K. E. et al. Directional patterns of cross frequency phase and amplitude coupling within the resting state mimic patterns of fMRI functional connectivity. Neuroimage 128, 238–251 (2016).

15. Cao, J. et al. Gastric stimulation drives fast BOLD responses of neural origin. Neuroimage 197, 200– 211 (2019).

16. Wolpert, N., Rebollo, I. & Tallon-Baudry, C. Electrogastrography for psychophysiological research: Practical considerations, analysis pipeline, and normative data in a large sample. Psychophysiology 57, e13599 (2020).

17. Glasser, M. F. et al. A multi-modal parcellation of human cerebral cortex. Nature 536, 171–178 (2016).

18. Li, Y. & Ngom, A. The non-negative matrix factorization toolbox for biological data mining. Source Code Biol. Med. 8, 10 (2013).

19. Cao, J., Wang, X., Chen, J., Zhang, N. & Liu, Z. The vagus nerve mediates the stomach-brain coherence in rats. Neuroimage 263, 119628 (2022).

20. Rebollo, I., Wolpert, N. & Tallon-Baudry, C. Brain–stomach coupling: Anatomy, functions, and future avenues of research. Curr. Opin. Biomed. Eng 18, 100270 (2021).

21. Cechetto, D. F. & Saper, C. B. Evidence for a viscerotopic sensory representation in the cortex and thalamus in the rat. J. Comp. Neurol. 262, 27–45 (1987).

22. Dum, R. P., Levinthal, D. J. & Strick, P. L. The spinothalamic system targets motor and sensory areas in the cerebral cortex of monkeys. J. Neurosci. 29, 14223–14235 (2009).

23. Simmons, W. K. et al. Keeping the body in mind: insula functional organization and functional connectivity integrate interoceptive, exteroceptive, and emotional awareness. Hum. Brain Mapp. 34, 2944–2958 (2013).

24. Rolls, E. T. The cingulate cortex and limbic systems for emotion, action, and memory. Brain Struct. Funct. 224, 3001–3018 (2019).

25. Capilla, A. et al. The natural frequencies of the resting human brain: An MEG-based atlas. Neuroimage 258, 119373 (2022).

26. Roy, M., Shohamy, D. & Wager, T. D. Ventromedial prefrontal-subcortical systems and the generation of affective meaning. Trends Cogn Sci (Regul Ed) 16, 147–156 (2012).

27. Ladabaum, U. et al. Gastric distention correlates with activation of multiple cortical and subcortical regions. Gastroenterology 120, 369–376 (2001).

28. van Oudenhove, L. et al. Cortical deactivations during gastric fundus distension in health: visceral pain-specific response or attenuation of “default mode” brain function? A H2 15O-PET study. Neurogastroenterol. Motil. 21, 259–271 (2009).

29. Mayeli, A. et al. Parieto-occipital ERP indicators of gut mechanosensation in humans. Nat. Commun. 14, 3398 (2023).

30. Levinthal, D. J. & Strick, P. L. Multiple areas of the cerebral cortex influence the stomach. Proc Natl Acad Sci USA 117, 13078–13083 (2020).

31. Das, A. et al. Spontaneous neuronal oscillations in the human insula are hierarchically organized traveling waves. eLife 11, (2022).

32. Sattler, N. J. & Wehr, M. Cortex-wide spatiotemporal motifs of theta oscillations are coupled to freely moving behavior. Front. Syst. Neurosci. 19, 1557096 (2025).

33. Ye, Q., Yan, D., Yao, M., Lou, W. & Peng, W. Hyperexcitability of Cortical Oscillations in Patients with Somatoform Pain Disorder: A Resting-State EEG Study. Neural Plast. 2019, 2687150 (2019).

34. Salmelin, R. & Hari, R. Characterization of spontaneous MEG rhythms in healthy adults. Electroencephalogr. Clin. Neurophysiol. 91, 237–248 (1994).

35. Yeo, B. T. T. et al. The organization of the human cerebral cortex estimated by intrinsic functional connectivity. J. Neurophysiol. 106, 1125–1165 (2011).

36. Engel, A. K., Gerloff, C., Hilgetag, C. C. & Nolte, G. Intrinsic coupling modes: multiscale interactions in ongoing brain activity. Neuron 80, 867–886 (2013).

37. Hoffstaedter, F. et al. The role of anterior midcingulate cortex in cognitive motor control: evidence from functional connectivity analyses. Hum. Brain Mapp. 35, 2741–2753 (2014).

38. Sauseng, P., Hoppe, J., Klimesch, W., Gerloff, C. & Hummel, F. C. Dissociation of sustained attention from central executive functions: local activity and interregional connectivity in the theta range. Eur. J. Neurosci. 25, 587–593 (2007).

39. Uddin, L. Q. Salience processing and insular cortical function and dysfunction. Nat. Rev. Neurosci. 16, 55–61 (2015).

40. Seeley, W. W. et al. Dissociable intrinsic connectivity networks for salience processing and executive control. J. Neurosci. 27, 2349–2356 (2007).

41. Andrews-Hanna, J. R., Smallwood, J. & Spreng, R. N. The default network and self-generated thought: component processes, dynamic control, and clinical relevance. Ann. N. Y. Acad. Sci. 1316, 29–52 (2014).

42. Chalas, N. et al. Respiration as a dynamic modulator of sensory sampling. Nat. Commun. 17, (2026).

43. Mahjoory, K., Schoffelen, J.-M., Keitel, A. & Gross, J. The frequency gradient of human resting-state brain oscillations follows cortical hierarchies. eLife 9, (2020).

44. Bastos, A. M. et al. Visual areas exert feedforward and feedback influences through distinct frequency channels. Neuron 85, 390–401 (2015).

45. Fries, P. Rhythms for Cognition: Communication through Coherence. Neuron 88, 220–235 (2015).

46. Michalareas, G. et al. Alpha-Beta and Gamma Rhythms Subserve Feedback and Feedforward Influences among Human Visual Cortical Areas. Neuron 89, 384–397 (2016).

47. Jensen, O. & Mazaheri, A. Shaping functional architecture by oscillatory alpha activity: gating by inhibition. Front. Hum. Neurosci. 4, 186 (2010).

48. Engel, A. K. & Fries, P. Beta-band oscillations--signalling the status quo? Curr. Opin. Neurobiol. 20, 156–165 (2010).

49. Dum, R. P., Levinthal, D. J. & Strick, P. L. Motor, cognitive, and affective areas of the cerebral cortex influence the adrenal medulla. Proc Natl Acad Sci USA 113, 9922–9927 (2016).

50. Voytek, B. & Knight, R. T. Dynamic network communication as a unifying neural basis for cognition, development, aging, and disease. Biol. Psychiatry 77, 1089–1097 (2015).

51. Vinck, M. et al. Principles of large-scale neural interactions. Neuron 111, 987–1002 (2023).

52. Bazinet, V., Vos de Wael, R., Hagmann, P., Bernhardt, B. C. & Misic, B. Multiscale communication in cortico-cortical networks. Neuroimage 243, 118546 (2021).

53. Vartiainen, J., Liljeström, M., Koskinen, M., Renvall, H. & Salmelin, R. Functional magnetic resonance imaging blood oxygenation level-dependent signal and magnetoencephalography evoked responses yield different neural functionality in reading. J. Neurosci. 31, 1048–1058 (2011).

54. Lankinen, K. et al. Consistency and similarity of MEG– and fMRI-signal time courses during movie viewing. Neuroimage 173, 361–369 (2018).

55. Hall, E. L., Robson, S. E., Morris, P. G. & Brookes, M. J. The relationship between MEG and fMRI. Neuroimage 102 **Pt** **1**, 80–91 (2014).

56. Tort, A. B. L., Komorowski, R., Eichenbaum, H. & Kopell, N. Measuring phase-amplitude coupling between neuronal oscillations of different frequencies. J. Neurophysiol. 104, 1195–1210 (2010).

57. Elam, M., Thorén, P. & Svensson, T. H. Locus coeruleus neurons and sympathetic nerves: activation by visceral afferents. Brain Res. 375, 117–125 (1986).

58. Teckentrup, V. et al. Non-invasive stimulation of vagal afferents reduces gastric frequency. Brain Stimulat. 13, 470–473 (2020).

59. Hong, G.-S. et al. Effect of transcutaneous vagus nerve stimulation on muscle activity in the gastrointestinal tract (transVaGa): a prospective clinical trial. Int. J. Colorectal Dis. 34, 417–422 (2019).

60. McCallum, R. W. et al. Mechanisms of symptomatic improvement after gastric electrical stimulation in gastroparetic patients. Neurogastroenterol. Motil. 22, 161–7, e50 (2010).

61. van Helden, D. F., Laver, D. R., Holdsworth, J. & Imtiaz, M. S. Generation and propagation of gastric slow waves. Clin. Exp. Pharmacol. Physiol. 37, 516–524 (2010).

62. Gharibans, A. A., Kim, S., Kunkel, D. & Coleman, T. P. High-Resolution Electrogastrogram: A Novel, Noninvasive Method for Determining Gastric Slow-Wave Direction and Speed. IEEE Trans Biomed Eng (2016) doi:10.1109/TBME.2016.2579310.

63. Tallon-Baudry, C., Loescher, M. & Clément, A. A dual-function framework of interoception: the information and coordination modes of interoceptive signaling. Trends Neurosci. 49, 161–172 (2026).

64. Fries, P. A mechanism for cognitive dynamics: neuronal communication through neuronal coherence. Trends Cogn Sci (Regul Ed) 9, 474–480 (2005).

65. Singer, W. Synchronization of cortical activity and its putative role in information processing and learning. Annu. Rev. Physiol. 55, 349–374 (1993).

66. Buzsáki, G., Logothetis, N. & Singer, W. Scaling brain size, keeping timing: evolutionary preservation of brain rhythms. Neuron 80, 751–764 (2013).

67. Rafael, O., et al. A cortical basis for perception of internal gut sensations. bioRxiv (2026).

68. Banellis, L., Rebollo, I., Nikolova, N. & Allen, M. Stomach–brain coupling indexes a dimensional signature of mental health. Nat. Mental Health (2025) doi:10.1038/s44220-025-00468-6.

69. Loescher, M., Wolpert, N. & Tallon-Baudry, C. Perceptual reaction times are coupled to the gastric electrical rhythm. bioRxiv (2026).

70. Khalsa, S. S., Berner, L. A. & Anderson, L. M. Gastrointestinal interoception in eating disorders: charting a new path. Curr. Psychiatry Rep. 24, 47–60 (2022).

71. Babo-Rebelo, M., Buot, A. & Tallon-Baudry, C. Neural responses to heartbeats distinguish self from other during imagination. Neuroimage 191, 10–20 (2019).

72. Kluger, D. S., Allen, M. G. & Gross, J. Brain-body states embody complex temporal dynamics. Trends Cogn Sci (Regul Ed) 28, 695–698 (2024).

73. Oostenveld, R., Fries, P., Maris, E. & Schoffelen, J.-M. FieldTrip: Open source software for advanced analysis of MEG, EEG, and invasive electrophysiological data. Comput. Intell. Neurosci. 2011, 156869 (2011).

74. Nolte, G. The magnetic lead field theorem in the quasi-static approximation and its use for magnetoencephalography forward calculation in realistic volume conductors. Phys. Med. Biol. 48, 3637–3652 (2003).

75. Van Veen, B. D., van Drongelen, W., Yuchtman, M. & Suzuki, A. Localization of brain electrical activity via linearly constrained minimum variance spatial filtering. IEEE Trans Biomed Eng 44, 867– 880 (1997).

76. Mitra, P. P. & Pesaran, B. Analysis of dynamic brain imaging data. Biophys. J. 76, 691–708 (1999).

77. Berens, P. circstat: a *MATLAB* toolbox for circular statistics. J. Stat. Softw. 31, (2009).

78. Unable to find information for 18371729.

79. Alexander-Bloch, A. F. et al. On testing for spatial correspondence between maps of human brain structure and function. Neuroimage 178, 540–551 (2018).

80. Chen, Y., Bressler, S. L. & Ding, M. Frequency decomposition of conditional Granger causality and application to multivariate neural field potential data. J. Neurosci. Methods 150, 228–237 (2006).

